# Experimental evidence that chronic outgroup conflict reduces reproductive success in a cooperatively breeding fish

**DOI:** 10.1101/2021.08.11.455992

**Authors:** Ines Braga Goncalves, Andrew N. Radford

## Abstract

Conflicts with conspecific outsiders are common in group-living species, from ants to primates, and are argued to be an important selective force in social evolution. However, whilst an extensive empirical literature exists on the behaviour exhibited during and immediately after interactions with rivals, only very few observational studies have considered the cumulative fitness consequences of outgroup conflict. Using a cooperatively breeding fish, the daffodil cichlid (*Neolamprologus pulcher*), we conducted the first experimental test of the effects of chronic outgroup conflict on reproductive investment and output. ‘Intruded’ groups received long-term simulated territorial intrusions by neighbours that generated consistent group-defence behaviour; matched ‘Control’ groups (each the same size and with the same neighbours as an Intruded group) received no intrusions in the same period. Intruded groups had longer inter-clutch intervals and produced eggs with less protein than Control groups. Despite this lower egg investment, Intruded groups provided more parental care, achieving similar hatching success to Control groups. Ultimately, however, Intruded groups had fewer and smaller surviving offspring than Control groups at 1-month post-hatching. We therefore provide experimental evidence that outgroup conflict can decrease fitness via cumulative effects on reproductive success, confirming the selective potential of this empirically neglected aspect of sociality.

## Introduction

In social species, conspecific outsiders often challenge groups and their members for resources and reproductive opportunities (1–5), and this outgroup conflict is theorised to be a driving force in the evolution of territoriality, social structure, group dynamics and cooperation (6–9). Empirical research in non-human animals has traditionally focused on aggressive outgroup contests, such as factors determining which individuals participate, their level of contribution and who wins (5,10), as well as the immediate fitness costs arising from loss of life, breeding position or territory (11,12). Recently, studies have begun to explore the effects of outgroup conflict beyond periods of active confrontation, documenting short-term behavioural changes (e.g. increased within-group affiliation) in the aftermath of single interactions with rivals (1-3,13). However, the cumulative build-up of threat posed by outsiders is also likely a potent stressor (14,15) and may disrupt within-group relationships (16,17), potentially generating long-term fitness consequences even in the absence of immediate direct effects from individual aggressive contests. Three observational studies have found associations between outgroup conflict and reproductive success, beyond the direct death of offspring during aggressive interactions: greater intergroup conflict during pregnancy was associated with improved foetal survival in both crested macaques (*Macaca nigra*) and banded mongooses (*Mungos mungo*) (4,18), but with lower birth rates and infant survival in chimpanzees (*Pan troglodytes verus*) (19). However, experiments are needed to test a causal link between outgroup conflict and reproductive success.

Here, we use the daffodil cichlid (*Neolamprologus pulcher*), a model species for the study of sociality, to test experimentally the reproductive consequences of chronically elevated outgroup conflict. *N. pulcher* is a highly territorial, cooperatively breeding fish species native to Lake Tanganyika. In the wild, groups comprising a breeding pair and 0–20 subordinates of both sexes (20) defend territories from predators, heterospecific competitors and conspecific intruders (21,22), with multiple small, contiguous territories often clustered together (23). Although individuals develop dear-enemy relationships with neighbours (24,25), aggressive disputes at shared borders are common (26) and intruding neighbours are attacked by all group members (27). Single intrusions are known to cause short-term changes in within-group behavioural interactions (2,28,29). Crucially, *N. pulcher* is a highly tractable experimental system—they are easily maintained in captive conditions, where groups display natural behaviour and breed regularly (20,30,31)—allowing controlled manipulations and detailed monitoring over extended periods.

## Results

To investigate the cumulative effect of elevated outgroup conflict on general reproductive behaviour, investment in eggs and parental care, and reproductive output, we simulated intrusions by neighbours at territorial borders in two long-term experiments (see Methods). In both experiments (which used different fish), 30 tanks with breeding shelters were arranged linearly in triplets: the end tanks (one for each of two treatments: 10 ‘Intruded’ and 10 ‘Control’ tanks per experiment) contained a dominant pair and a subordinate; the middle tank contained a dominant pair who comprised the shared neighbours used as intruders (Fig 1A). Within each triplet, the same-sex dominants in all tanks, and the subordinates in experimental tanks (i.e. the end tanks), were size-matched. ‘Intruded’ groups experienced regular territorial intrusions by neighbouring individuals (mean±s.e. intrusions per week, Experiment I: 4.8±0.1; Experiment II: 3.9±0.3); ‘Control’ groups did not receive intrusions. In Experiment I, we allowed each group to raise all clutches produced over a 13-week period; in Experiment II, we collected the first clutch produced (Fig 1B). Intruded groups in Experiment I perceived the presence of neighbours in their territory as intruders, exhibiting 4.5 times more defensive actions than Control groups at the start of the study (Wilcoxon signed-test: V=0, n=20, p=0.002; Fig 2A); this difference persisted until the end of Experiment I (4.7 times more defence; paired t-test: t_9_=4.33, p=0.002; Fig 2A). Within the Intruded treatment, dominant individuals, but not subordinates, increased their defensive efforts between the start and the end of the study (dominant male: t_9_=3.50, p=0.007; dominant female: V=4.5, n=20, p=0.022; subordinate: V=5.0, n=12, p=1.0; Fig 2B); this pattern was not driven by changes in Intruder responsiveness to the focal group (t_9_=0.55, p=0.594). Greater defensive efforts by dominant individuals at the end of the study may be a by-product of age or size-mediated behavioural changes—an effect of size on defensive efforts has been documented in at least dominant males in this species (32)—or may have been mediated by the presence of offspring that need safeguarding from intruders (33).

**Figure 1.**
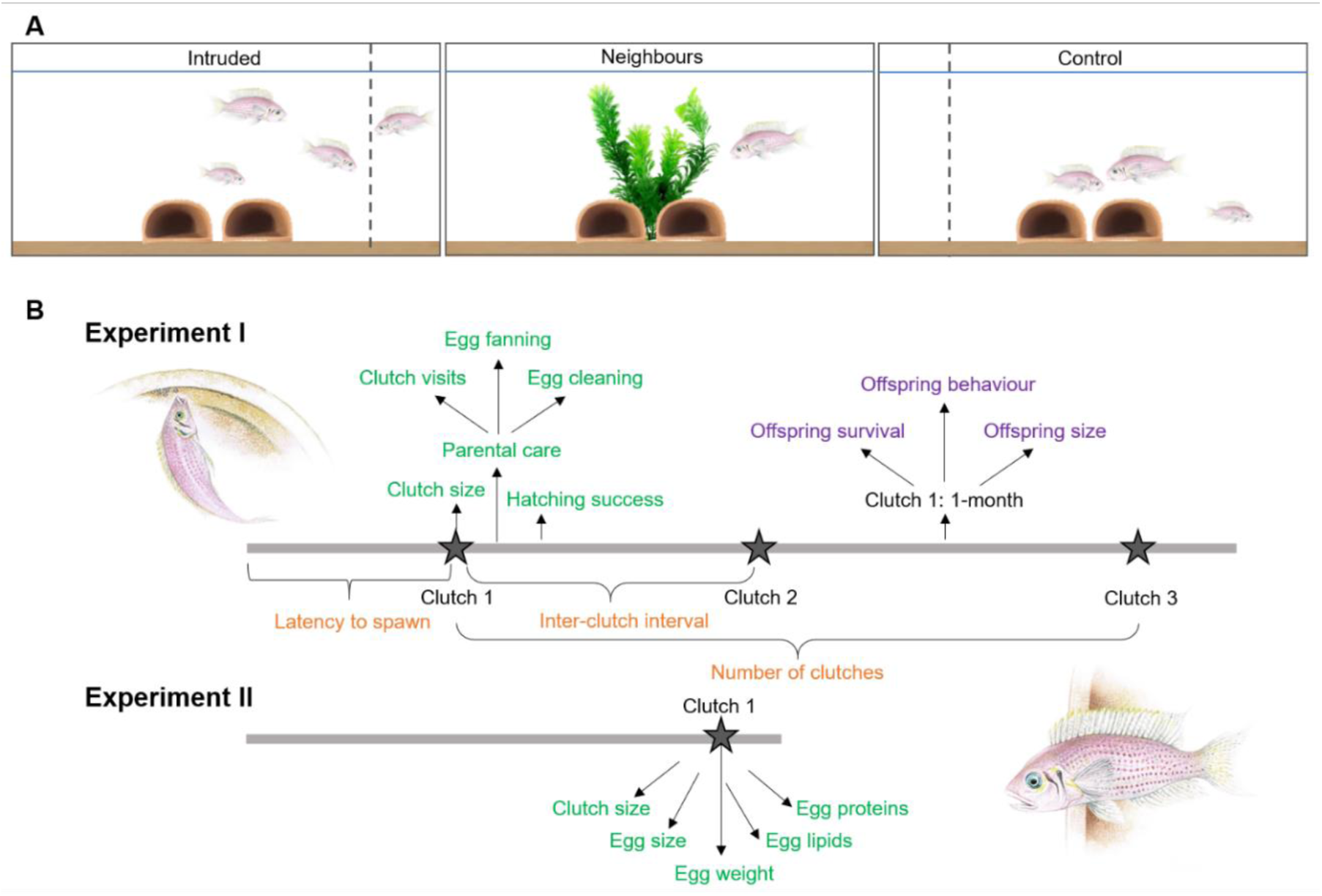
Experimental set-up and timelines. **(A)** Experimental set-up showing a linearly arranged tank triplet, with size-matched experimental groups (Intruded and Control) at the ends and the tank with the neighbour pair at the centre. The female neighbour is presented in the Intruded group tank to illustrate an intrusion along the border of the resident group’s territory. **(B)** Timelines show measures of general reproductive behaviour (orange), egg and parental-care investment (green), and reproductive output (purple).

**Figure 2.**
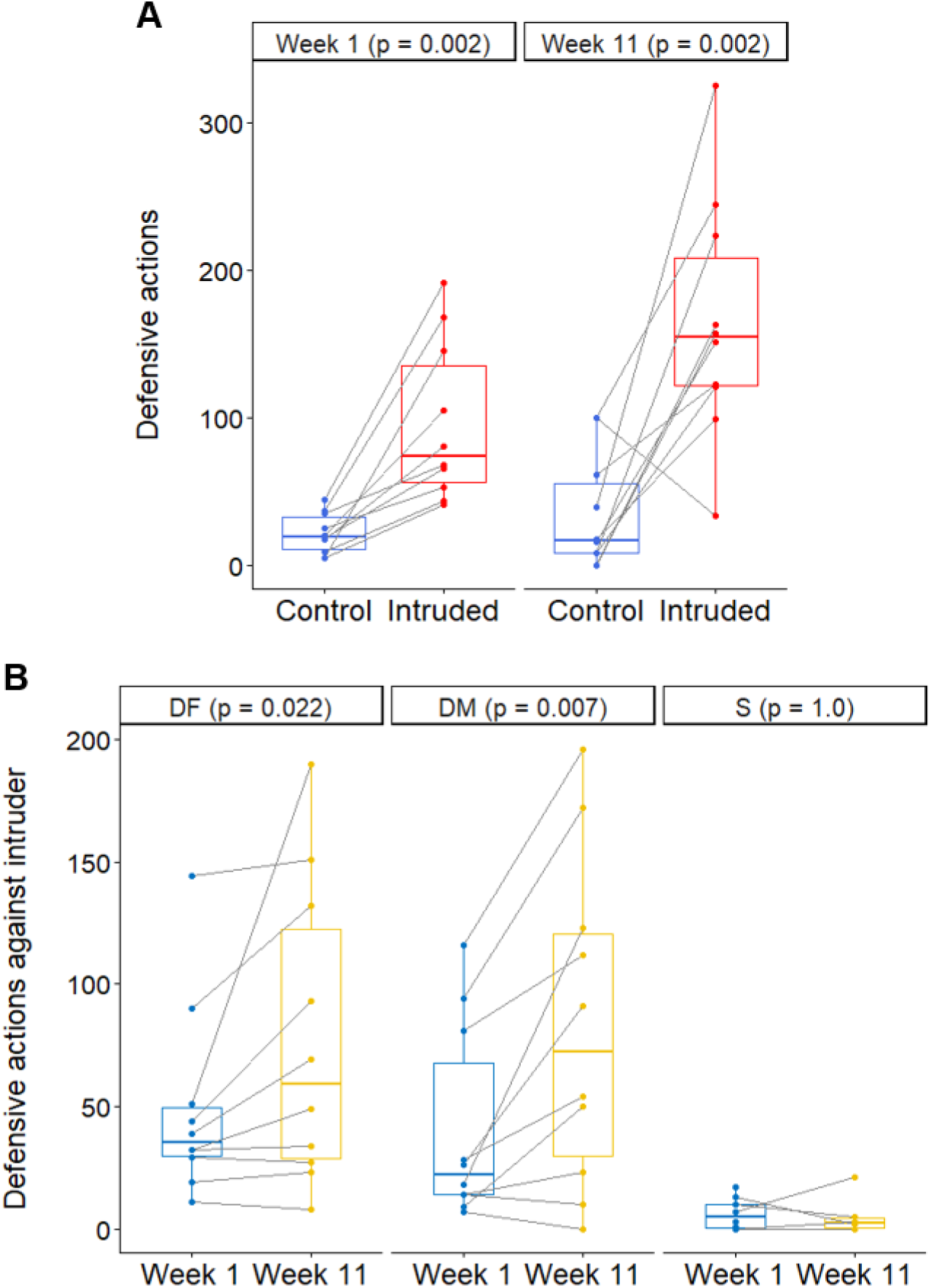
Defensive actions. **(A)** Group defensive actions displayed per 10 min trial, towards a transparent partition (blue) or the intruding female neighbour (red) in weeks 1 and 11 of Experiment I (N=20 groups). **(B)** Number of defensive actions displayed by the dominant female (DF, N=10), dominant male (DM, N=10) and subordinate (S, N=6) towards the intruding female neighbour during weeks 1 (blue) and 11 (yellow) of Experiment I. Boxplots show medians, 25% and 75% quartiles, and whiskers representing 95% confidence intervals; dots are raw data, with lines connecting matched groups (A) or repeated measures on individuals (B).

### General reproductive behaviour

When faced with prolonged challenging conditions, organisms mount a stress response that aims to maximise survival, potentially at the cost of other functions such as reproduction (34). In our Experiment I, repeated territorial intrusions did not significantly affect the likelihood of spawning (McNemar test: *χ*^2^_1_=0.36, p=0.547), the latency to first spawn (linear mixed model (LMM): *χ*^2^_1_=1.07, p=0.301; S1a Table) or the number of clutches produced (Wilcoxon rank-sum test: V=15, N=10, p=0.396). However, Intruded groups had inter-clutch intervals that were 40% longer than Control groups (*χ*^2^_1_=3.89, p=0.049; Fig 3A), after controlling for a positive effect of female size (S1b Table). Our finding aligns with previous studies showing that other chronic stressors negatively impact fish reproductive rates (35,36). Longer inter-clutch intervals likely result in fewer breeding attempts per season, which is why inter-birth interval is a life-history trait commonly used to assess female reproductive success across taxa (37).

**Figure 3.**
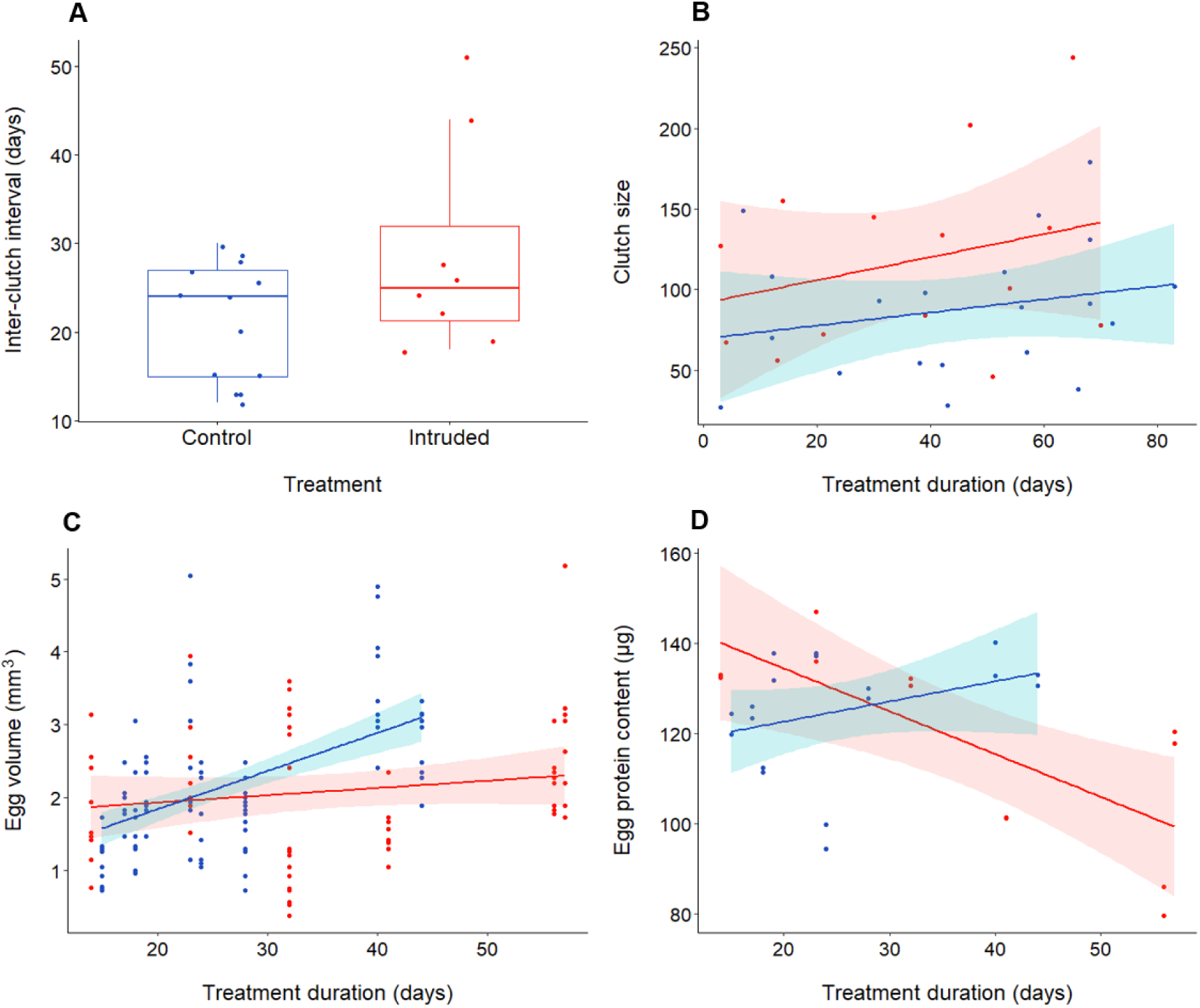
Effects of chronically elevated outgroup threat on egg investment. Effects of chronically elevated outgroup threat (red) relative to control conditions (blue) on: **(A)** inter-clutch interval (N=21 intervals); **(B)** clutch size (N=34 clutches); **(C)** egg volume (N=15 clutches); and **(D)** egg protein content (N=15 clutches). Boxplot shows median and 25% and 75% quartiles, with whiskers representing 95% confidence intervals; scatterplots display lines of best fit and associated confidence intervals; dots are raw data.

### Egg and parental-care investment

In fish, females can modify egg number, size and nutritional content in response to the prevailing ecological and social conditions (38,39), with trade-offs between egg number and quality apparent in stressful situations (39). Where our cichlid groups were allowed to produce just one clutch (Experiment II), clutch size was not significantly affected by treatment (ANCOVA: F_1,12_=0.15, p=0.860) or its interaction with treatment duration (F_1,10_=0.05, p=0.835), after controlling for a positive effect of female size (F_1,14_=11.34, p=0.005). However, where groups could produce multiple clutches throughout the 13-week study period (Experiment I), there was a near-significant effect of the interaction between treatment and treatment duration on clutch size: over time, Intruded females showed a tendency to produce larger clutches relative to Control females (LMM, parameter estimate [PE]=0.85, 95% confidence interval [CI]=-0.04–1.70, *χ*^2^_1_=3.55, p=0.059; S2 Table; Fig 3B). Using clutches from Experiment II, we assessed how increased outgroup conflict affected egg size and nutrient allocation. Whilst there was no significant effect of treatment on egg weight (*χ*^2^_1_=0.47, p=0.495; S3a Table), there was a significant effect of the interaction between treatment and treatment duration on egg volume (*χ*^2^_1_=4.34, p=0.037; S3b Table; Fig 3C): females who took longer to spawn produced eggs with larger volume in the Control treatment (PE=0.05, 95% CI=0.02–0.09, t_7.14_=2.95, p=0.021) but not in the Intruded treatment (PE=0.009, 95% CI=-0.019–0.037, t_4.13_=0.59, p=0.584). There was no significant treatment difference in egg lipid content (*χ*^2^_1_=0.03, p=0.856; table S3c), but egg protein content was significantly affected by the interaction between treatment and treatment duration (*χ*^2^_1_=7.65, p=0.006; S3d Table; Fig 3D): mean egg protein content showed a tendency to decrease with treatment duration in the Intruded treatment (PE=-0.95, 95% CI=-1.75–-0.15, t_4.00_=-2.25, p=0.088), but did not change in the Control treatment (PE=0.45, 95% CI=-0.43–1.33, t_7.00_=0.98, p=0.361). Overall, chronic outgroup conflict affected reproductive investment: Intruded females tended to invest in larger clutches the longer the treatment continued, but eggs in those later clutches contained less protein and, unlike in the Control groups, were not larger in size. Our results are in line with previous work on *N. pulcher* showing flexible adjustment of egg size in response to the social environment (40) and to repeated acute handling stress (35). Reductions in egg protein content, but not in lipid content, by females exposed experimentally to elevated glucocorticoids during egg production, have also been reported in eastern fence lizards, *Sceloporus undulatus* (41). Greater investment into egg number over quality is expected in more variable environments or when the probability of future reproduction is low (42).

Parental care extends the opportunity for adjustment of reproductive investment in response to the prevailing conditions, allowing compensation for lower offspring quality (42). In Experiment I, repeated territorial intrusions had an increasingly positive effect on the number of parental-care events (S4 Table) and the time spent in such activities (S5 Table); the former are detailed here. The number of clutch visits was significantly affected by the interaction between treatment and treatment duration (LMM: *χ*^2^_1_=4.85, p=0.028; S4a Table; Fig 4A): clutch visits increased over time in the Intruder treatment (PE=0.29, 95% CI=0.05–0.57, t_10.08_=2.29, p=0.045) but not in the Control treatment (PE=-0.03, 95% CI=-0.21–0.15, t_17.00_=-0.34, p=0.737). Similarly, treatment duration affected the number of care behaviours performed in a treatment-dependent manner (interaction: *χ*^2^_1_=4.49, p=0.034; S4b Table; Fig 4B): caring behaviour increased significantly over time in the Intruded treatment (PE=0.44, 95% CI=0.17–0.72, t_10.92_=3.22, p=0.008) but not in the Control treatment (PE=0.09, 95% CI=-0.10–0.28, t_14.14_=0.92, p=0.374). This pattern was driven by a significant interaction between treatment and treatment duration in the number of egg-cleaning events (*χ*^2^_1_=4.47, p=0.035; S4c Table)—with egg-cleaning events increasing more over time in the Intruded treatment (Intruded: PE=0.26 95% CI=0.10–0.42, t_10.03_=3.35, p=0.007; Control: PE=0.09, 95% CI=0.005–0.180, t_12.71_=2.14, p=0.052)—and a trend in the same direction for the number of egg-fanning events (*χ*^2^_1_=2.90, p=0.089; S4d Table). Our results contrast those of studies that have shown reductions in parental (43,44) and alloparental (45) care in response to immediate or short-term stressful situations but, as effects became evident over the course of the experiment, it is possible that acute and chronic stressors elicit opposing behavioural responses. The increased parental care in the Intruded treatment may have compensated for lower relative egg investment (see above) because there was no significant treatment difference in hatching success (*χ*^2^_1_=2.39, p=0.123; S6 Table; Fig 4C). Similar hatching success does not, however, necessarily equate to similar offspring quality (46).

**Figure 4.**
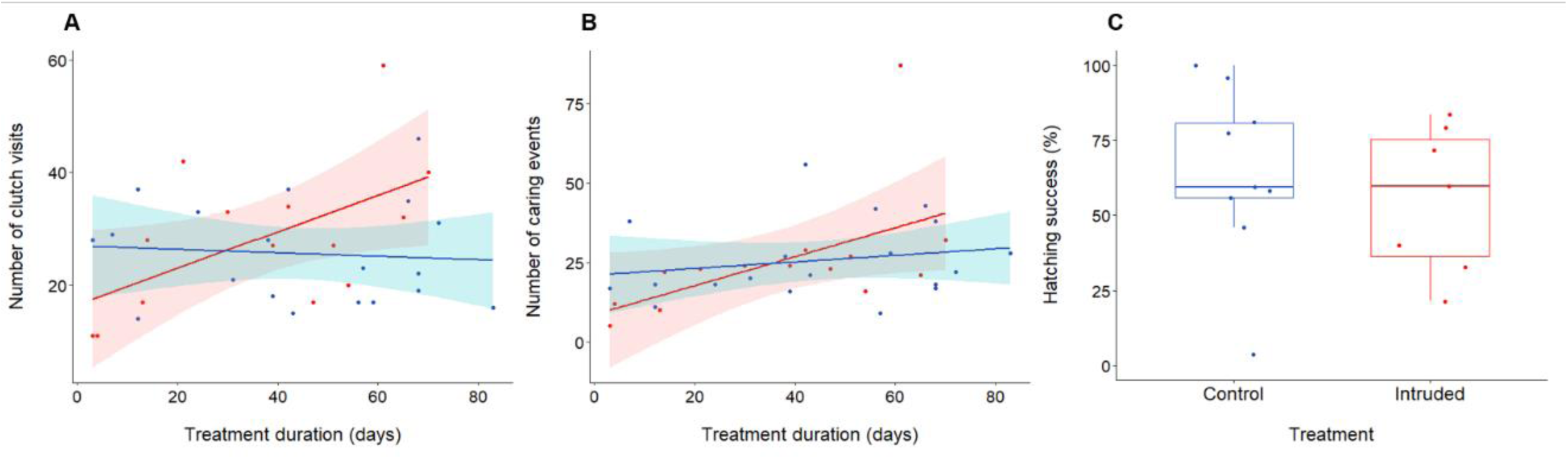
Effect of chronically elevated outgroup threat on parental-care investment and hatching success. Effects of chronically elevated outgroup threat (red) relative to control conditions (blue) on: **(A)** number of clutch visits (N=33 clutches); **(B)** number of clutch caring (cleaning and fanning) events per 10 min observation (N=33 clutches); and **(C)** offspring hatching success (N=16 clutches). Scatterplots display lines of best fit and associated confidence intervals; boxplot shows median and 25% and 75% quartiles, with whiskers representing 95% confidence intervals; dots are raw data.

### Reproductive output

Overall, the absolute number of offspring surviving to 1 month decreased significantly over treatment duration in the Intruded treatment relative to the Control treatment (LMM, interaction: *χ*^2^_1_=6.58, p=0.010; S7a Table; Fig 5A). This result was driven by a tendency for fewer offspring to survive to 1 month of age, over time, in the Intruder treatment (PE=-0.43, 95% CI=-0.900–-0.008, t_7.42_=3.35, p=0.083) but for the reverse pattern in the Control treatment (PE=0.49, 95% CI=-0.06–1.05, t_12.00_=-2.00, p=0.110). Fewer offspring were alive at 1 month not because of a significant difference in relative offspring survival (%) from hatching (*χ*^2^_1_=1.53, p=0.216; S7b Table) but because, in contrast to the Control treatment, the number of hatched fry in the Intruded treatment did not positively affect number of offspring that reached 1 month of age (interaction: *χ*^2^_1_=5.12, p=0.024; Intruded: PE=-0.17, 95% CI=-0.36–0.02, t_2.01_=-1.77, p=0.218, Control: PE=0.90, 95% CI=0.52–1.32, t_4.31_=4.86, p=0.009; S7c Table). Low offspring survival may result from stress effects on parents that reduce the quality of their offspring, the direct early-life experience of outgroup conflict on offspring or a combination of both (47,48). Alternatively, adults in the Intruded treatment may have displayed stress-induced filial cannibalism; social perturbations have been shown to induce egg-cannibalism by males (30), though we are not aware of reports of within-group fry or juvenile cannibalism in the study species.

**Figure 5.**
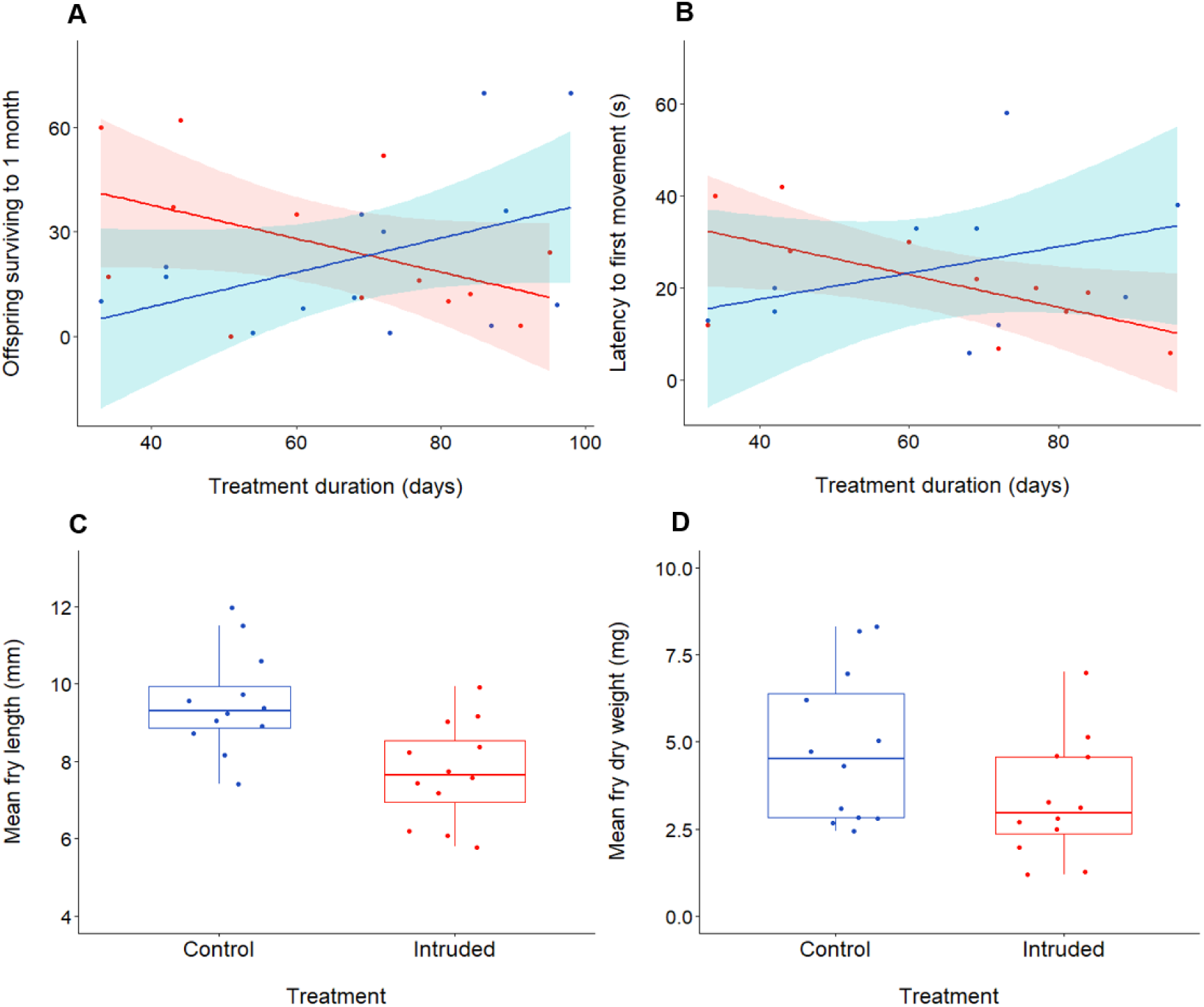
Effect of chronically elevated outgroup threat on reproductive output. Effect of chronically elevated outgroup threat (red) compared to control conditions (blue) on: **(A)** number of offspring surviving to 1 month (N=27 clutches); **(B)** latency to first movement of offspring post-stimulus (N=21 clutches); **(C)** mean offspring standard length (N=24 clutches); and **(D)** mean offspring dry weight (N=24 clutches). Scatterplots display lines of best fit and associated confidence intervals; boxplots show median and 25% and 75% quartiles, with whiskers representing 95% confidence intervals; dots are raw data.

Maternal and early-life stress can also influence a suite of offspring behaviours, including activity levels, shoaling decisions, and anti-predator responses (49–51). Normally, on detecting danger, young *N. pulcher* swiftly sink to the bottom and remain motionless to blend with the background environment (23,52). To investigate offspring behaviour, we placed 5–10 1-month-old young per clutch in a novel environment (20×20 cm container). Following a 30-min settling period, there was no significant treatment difference in offspring baseline activity level (proportion of active individuals, LMM: *χ*^2^_1_=0.95, p=0.329; S8a Table) or mean nearest-neighbour distance (*χ*^2^_1_=0.13, p=0.714; S8b Table). After this undisturbed period, we assessed the response to a startle stimulus. Treatment had no significant effect on offspring latency to freeze immediately post-stimulus (*χ*^2^_1_=0.03, p=0.855; S8c Table). However, after controlling for a positive effect of offspring size and a negative effect of pre-stimulus activity level, latency for the first offspring to move post-stimulus was significantly affected by the interaction between treatment and treatment duration (*χ*^2^_1_=4.82, p=0.028; S8d Table; Fig 5B): post-stimulus latency to move decreased significantly over time in the Intruded treatment (PE=-0.35, 95% CI=-0.64–-0.07, t_9.00_=-2.40, p=0.040) but not in the Control treatment (PE=0.29, 95% CI=-0.20–0.78, t_8.00_=1.13, p=0.290). The speed of return to activity is thought to determine vulnerability to predation, because prey movement enhances detection and attack rates by predators (51,53).

In fish, offspring size is influenced by initial egg size and parental-care quality, as well as by parental stress and early-life experience (38,47,54). Immediately following the behavioural tests, we sacrificed the offspring to assess standard length and dry weight. Compared to Control individuals, Intruded offspring were 19% shorter (LMM: *χ*^2^_1_=4.89, p=0.027; Fig 5C), after controlling for a negative effect of treatment duration (S9a Table), and were 30% lighter (*χ*^2^_1_=3.15, p=0.076; S9b Table; Fig 5D). The smaller size at 1-month of age indicates that outgroup conflict hinders early-life growth, despite unlimited access to food in our experiment. Size in early life is a key determinant of survival (55) and reproductive success (56) in fishes and, in the daffodil cichlid in particular, adult body size is a key determinant of lifetime reproductive success because it correlates positively with fecundity in females and with dominance acquisition in both sexes (20).

## Discussion

Our experimental results show that chronic elevation of outgroup conflict negatively impacted adult breeding timings and investment in egg quality; whilst there was a compensatory positive influence on parental care, an increased outgroup threat ultimately resulted in fewer offspring of smaller size surviving to 1-month of age. Our findings are in-line with previous correlational work showing reductions in chimpanzee reproductive rate and offspring survival associated with greater levels of intergroup threat (19). Tests of causal links between outgroup conflict and reproductive success have previously been lacking due to the inherent difficulties of carrying out long-term manipulations that allow cumulative fitness impacts to be assessed. We overcame this by using a model fish species that enables both extended manipulations in controlled conditions and the collection of behavioural and reproductive data (20). In doing so, we demonstrated that chronic outgroup conflict negatively impacts the fitness of multiple generations: adults suffer reductions in current reproductive output, whilst the lower quality of surviving offspring means that they are potentially less likely to achieve dominance and thus direct reproductive success later in life.

We uncovered outgroup-conflict effects in several aspects of reproductive investment and output that became stronger the longer the experimental treatment continued. These findings highlight that, to ascertain the full fitness consequences in social animals, it is important to consider both direct and indirect effects and to assess cumulative impacts in addition to those arising from single contests. Since outgroup conflict can be a potent social stressor, capable of stimulating acute and chronic stress responses (14), stress is a likely mechanism underpinning the reproductive consequences seen. Studies examining natural stress responses (57,58), and experimental manipulations of glucocorticoids (38,41,48), have demonstrated how stress can affect adult reproductive measures in ways similar to those uncovered in our work. Likewise, early-life stressors can have long-lasting effects on offspring phenotypes and fitness (47,59). Some of the fitness costs documented here may also have been indirectly caused by outgroup conflict-mediated disruptions to within-group social relationships. First, within-group aggression is promoted by the mere presence of neighbours in complex time-, rank- and sex-dependent ways (16) and may be further dependent on the extent of interactions between individual residents and neighbours (60). Second, territorial intrusions cause (at least) short-term changes to within-group social interactions (2,29). Cumulative effects of outgroup conflict are likely to have a wide influence, beyond reproductive success, and thus deserve further research attention moving forward.

There are obvious trade-offs in conducting captive vs field studies. In the context of cumulative outgroup-conflict consequences, long-term experimental manipulations are logistically and ethically challenging, which is why the few previous studies have reported only correlational data (4,18,19). Our captive experiments allowed for tight control of the environment and close monitoring of a variety of response measures, but consideration is needed of the ecological validity of the intrusion regime. In the wild, *N. pulcher* groups are often surrounded by several adjacent groups (23), so that territories are intruded by neighbours, as well as outsiders from further afield, predators and heterospecific competitors, from any direction. Therefore, though short in duration, intrusions likely take place several times daily, requiring continuous vigilance and quick behavioural responses, risking potential physical injury and leaving residents more vulnerable to predation (26,27,61). Our simulated intrusions by one of two neighbouring individuals, at the edge of the territory, occurred less that once daily. Our set-up also precluded individuals from injuring each other, the intruder from committing infanticide, sneaking a breeding attempt or taking-over part or all the territory, and individuals from having to trade-off foraging time against vigilance efforts (due to plentiful food availability), all in the absence of predation. Thus, relative to wild conditions, our lab intrusions were likely longer but less frequent, and posed relatively low threat to focal groups within an overall less challenging environment, increasing chronically the perception of outgroup threat that allowed us to assess cumulative indirect effects of outgroup conflict (beyond injury, death and infanticide) without continuously and acutely stressing our focal subjects. Accordingly, unlike in previous work on the effects of repeated acute stressors on reproduction in this species (35), our treatments did not negatively impact adult body mass or egg mass, had no impact on latency to breed and a much smaller impact on inter-clutch interval durations (40% vs >200% increase in duration); experimental neighbours (i.e. intruder pairs) had the same likelihood of spawning as Intruded groups (six of 10 pairs spawned), no different from Control groups. Ultimately, future field experiments are needed to confirm the causal impact of chronic outgroup conflict on reproductive success, but many challenges will need to be overcome first.

We demonstrate that chronically increased outgroup conflict can reduce reproductive success, even in the absence of physical fights with rivals that may also result in injury, death or loss of territory or breeding position (4,11,12), or cause offspring death (4,33). Outgroup conflict can clearly, therefore, influence fitness in myriad ways, lending support to the theory that it is likely a powerful selective force in social evolution with respect to, for example, group dynamics, social structure and cooperation (6–8). Given the widespread taxonomic occurrence of outgroup conflict, we advocate further experimental testing of what is arguably the most neglected aspect of sociality.

## Materials and Methods

### Fish husbandry

We used a captive population of daffodil cichlids, *Neolamprologus pulcher*, at the University of Bristol; work was approved by the University of Bristol Ethical Committee (University Investigator Number: UB/16/049 + UB/19/059). All fish groups were housed in 70-L tanks (width × length × height: 30 × 61 × 38 cm) that formed their territory (as in 2). Each tank contained 2–3 cm of sand (Sansibar river sand), a 75-W heater (Eheim), a filter (Eheim Ecco pro 130), a thermometer (Eheim), two flowerpot halves (10 cm wide) that served as breeding shelters at the centre of the territory, an artificial plant and a small tube hanging close to the water surface to provide extra shelter for the subordinate in focal groups. Fish were fed twice daily: alternating between frozen brine shrimp, water fleas, prawns, mosquito larvae, mysid shrimp, bloodworms, cichlid diet, spirulina, copepods and krill in the mornings from Monday to Friday; and dry fish flakes in the evenings and weekends. Water temperature was maintained constant (mean±se, Experiment I: 26.9±0.08°C, Experiment II: 26.7±0.10°C) and room lights were set on a 13L:11D hour cycle (daylight from 7 am to 8 pm). Water quality tests (pH, nitrates, nitrites, conductivity and ammonia) and 10% water changes were performed weekly to maintain water quality levels.

### Experimental set-up

In both experiments, we organised tanks in triplets (N=10), positioned in a line with the short sides about 0.5 cm apart, so that neighbours could always see each other. In each triplet, the central tank housed a breeding pair that was a common neighbour to the two focal groups (each comprising a breeding pair and an adult helper) on either side. Groups of three, although at the low range of natural group sizes for this species, are common in nature (26) and frequently used in laboratory studies (2,35,62–64). We formed groups separately (with different fish used for the two experiments) using standard procedures (2) approximately two weeks prior to the start of each experiment. In Experiment I, the 20 focal groups included 12 with female helpers and eight with male helpers; within a triplet experimental groups had same-sex helpers. In Experiment II, the 20 focal groups all had female helpers. Female helpers were favoured because although all subordinates face increasing risks of eviction as they grow (28), males are more likely to parasitise reproductive events resulting in eviction from the group (65) rendering groups less stable, and group size has been shown to affect breeder egg parameters (40). To minimise same-sex aggression and to aid individual identification, we ensured that each dominant was at least 5 mm larger than the same-sex subordinate in the focal group (mean±se size difference, Experiment I: 10.7±1.0 mm; Experiment II: 22.4±1.3 mm). To minimise potential size-difference effects between treatments, and control for the influence of female size on reproductive measures (66), we size-matched (in standard length) breeders in all tanks in a triplet to each other. In Experiment I, focal-group dominant males were 55.2±1.3 mm, dominant females were 53.4±0.9 mm, subordinate males were 41.8±0.9 mm and subordinate females were 44.3±1.3 mm long, while males were 59.3±2.6 mm and females were 54.8±1.6 mm long in the middle-tank breeding pairs. In Experiment II, focal-group dominant males were 73.1±1.3 mm, dominant females were 64.4±0.8 mm and subordinate females were 41.8±1.0 mm long, while males were 71.4±2.7 mm and females were 63.6±1.8 mm long in the middle-tank breeding pairs. In Experiment I, focal individuals in the two treatments remained size-matched at the end of the study in standard length (mm, paired t-test, DM: t_9_=0.10, p = 0.920; DF: t_9_ = 1.67, p = 0.130; Wilcoxon signed rank test, Sub: V = 15.5, n= 12, p = 0.343) and in body mass (g, paired t-test, DM: t_9_=0.64, p = 0.540; DF: t_9_ = 0.11, p = 0.914; Wilcoxon signed rank test, Sub: V = 18, n= 12, p = 0.156). Focal individuals were not re-measured at the end of Experiment II because groups finished treatment at different times (see Figure 1).

For each triplet, we randomly allocated (by flip of a coin) the side tanks to one of two experimental treatments, Control and Intruded; the central tank provided the intruders used in the Intruded treatment. Experiment I spanned 13 weeks to allow for multiple spawning bouts; we destroyed clutches produced before groups had experienced at least three intrusions (Control=1 clutch, Intruded=2 clutches) as groups would likely not have been significantly affected by treatments within such a short timeframe. In Experiment II, following eight intrusions during the first 13 days, the first clutch produced by each group marked the end of treatment for that group; two weeks represents about half the time of a regular breeding cycle (30,67). Twelve clutches were destroyed during the initial 13 days of treatment (Control=8 clutches, Intruded=4 clutches). Experiment II was run for 11 weeks to maximise the number of focal groups that produced a clutch.

### Simulated territorial intrusions

We selected the side of the focal Intruded tank where each intrusion took place (near or far from the neighbour tank) and the identity of the intruder (male or female neighbour) pseudo-randomly: by flip of a coin, but no more than four of the same side or sex in a row. In the wild, dominant individuals of both sexes undertake regular forays to nearby territories (68) and may, thus, return to their own territories from any direction. At the start of an intrusion (or equivalent in the Control tanks), we slid down one transparent and one opaque flexible partition (0.75 mm white ViPrint) through single-channel PVC tracks glued to the long walls, 8 cm from the tank edge, creating a side compartment at the edge of the territory of the focal group (2). Then, we netted out the pre-selected neighbour and placed it in the side compartment of the focal tank, obscured from view of the resident group for a 5-min settling period. Brief handling experiences do not affect behaviour adversely in this species (2,69). After the settling period, we removed the opaque partition in the focal tank to reveal the intruder to the resident group for 10 min, a trial duration that falls comfortably with the range (5–20 min) often used in laboratory and field studies in this species (22,68,70). At the end of the intrusion period, we replaced the opaque partition in the focal tank and placed another opaque partition between the focal and neighbour tanks, netted the intruder and transferred it back to the neighbour tank out of sight of the focal group. Concurrently, in the matching Control group, we conducted the same sequence of placement and removal of opaque and transparent partitions as in the Intruded tank, but in the absence of an intruder. To minimise the impact of human presence on the behaviour of the experimental subjects, the experimenter hid behind a curtain during the period that the intruder was visible to the resident group.

In Experiment I, we filmed (Sony Handycam HDR-XR520) two simulated intrusions, one at the start (Week 1) and another towards the end (Week 11) of the study, to assess how focal groups responded to the presence of a neighbour in their territory, and how their response changed over time. For consistency, all filmed intrusions had the female neighbour as intruder placed on the side of the tank closest to the neighbour tank. Similarly, we filmed the behavioural responses of the Control groups to the presence of the transparent partition in their territory. Subordinate sample size was smaller (N=12) at the end of the study due to deaths unrelated to the experiment (fish jumped out of tank) and evictions (Control=4, Intruded=4). Following previously established protocols for this species (71,72), we recorded using JWatcher (v1.0; Macquarie University, Sydney, Australia) the frequencies of aggressive behaviours, including attacks (rams and bites) and aggressive displays (aggressive postures, frontal displays and fast approaches), directed at the intruder or the transparent partition by each group member. From the Intruded videos, we also recorded Intruder responsiveness towards the focal group, by assessing proportion of time active and facing the focal group, as in (2).

### General reproductive behaviour

To assess the impact of outgroup conflict on the likelihood of spawning, latency to spawn, number of clutches produced and mean inter-clutch interval (Experiment I), we scanned all focal tanks every day for new clutches.

### Investment in eggs and parental care

To evaluate the impact of outgroup threat on clutch size, we photographed all clutches (Experiment I: N=34, Experiment II: N=15) the day after they were laid and counted the numbers of eggs using ImageJ (version 1.46r, National Institutes of Health, USA). Hatching success could only be assessed in a subset of clutches (N=16) because the hatchlings of several clutches took cover on the sand or on the plant and could not be counted reliably. From each clutch in Experiment II, we separated: (a) 10–12 eggs for assessment of egg size; (b) two samples of 6–11 eggs for lipid analysis; and c) two samples of five eggs for protein analysis. Two clutches (1 Control, 1 Intruded) were laid over two days; we collected two samples for egg-size assessment from these clutches. Samples (b) and (c) were stored at -20°C until extraction.

Using the egg-size sample, we measured the length and width of each egg with a stereo microscope (2x magnification) and a graticule. We then calculated the effective diameter of each egg (i.e. the diameter if it were perfectly round) as the cube root of the length multiplied by the square of the width, and used the effective diameter to calculate the volume assuming a spherical shape (73). We used all egg volumes from each sample in the analysis. After the eggs were individually measured, all eggs from the same clutch were placed together in a petri dish with wax paper and dried in a heating cupboard at 70°C for 48 h, weighed twice (ME5, Sartorius, Göttingen, Germany), returned to the heating cupboard for another 24 h and weighed twice again; all four measurements were used in the analysis.

To assess the total lipid content of eggs, we used the colorimetric sulfo-phosphato-vanillin (SPV) method for microquantities (74). Briefly, we dried the samples at 70°C for 24 h and weighed them twice to the nearest microgram (CPA26P, Sartorius, Göttingen, Germany). We transferred the samples into 15 ml round-bottom glass test tubes (16×150 mm) and crushed them with a glass rod before adding 10 ml of chloroform-methanol solution (1:1 v/v). We extracted 0.5 ml of supernatant from each sample into new test tubes and placed them in a dry bath (LSE single block digital; Corning Ltd, Barry, UK) at 100°C for 15 min to evaporate the solvent. After, we added 0.2 ml of sulphuric acid to the samples and placed them in the dry bath for a further 10 min. Once the samples had cooled to room temperature, we added 4.8 ml of vanillin-ortho-phosphoric acid 85% reagent (1.2 g/l), thoroughly mixed the samples for 30 s with a vortex mixer (Bibby Scientific, Stone, UK), and transferred 1 ml of the solution into 1 ml polystyrene semi micro cuvettes. Using a spectrophotometer (WPA Biowave UV/Vis; Biochrom Ltd, Cambridge, Uk) at 525 nm, calibrated with a vanillin/phosphoric acid only blank, we took three absorbance readings from each sample and calculated their mean. The mean sample absorbance values were plotted against a standard curve to extrapolate their total lipid content (µg). The standard curve was prepared using eight serial dilutions of analytical soybean oil solution (0.917g/ml, Sigma Aldrich) in methanol: chloroform (1:1).

To assess the total protein content of eggs, we used the Bradford method (75) following specifications by Alasmari and Wall (76). The egg samples were transferred into 15 ml round-bottom borosilicate glass test tubes (16×150 mm) and crushed with a glass rod. We added 0.5 ml of phosphate buffer (100 mM of monopotassium phosphate (KH2PO4, 1 mM of ethylenediaminetetraacetic acid (EDTA) and 1 mM of dithiothreitol (DTT) dissolved in distilled water, mixed with an aqueous solution of potassium phosphate dibasic (K2HPO4) to achieve pH=7.4) to dissolve the sample, followed by another 0.5 ml of buffer to clean the glass rod. After, we thoroughly mixed the samples for 30 s with a vortex mixer (Bibby Scientific, Stone, UK), we transferred 0.1 ml into new tubes, added 2.9 ml of buffer, and vortexed the new solutions for another 30 s to perform 1:30 dilutions. From these dilutions, we transferred 1 ml of the solution into a new tube, added 1 ml of Bradford reagent and vortexed the resulting solution for 1 min. After waiting 5 min at room temperature, we transferred 1 ml of the solution into 1 ml polystyrene semi micro cuvettes and read the absorbance values on a spectrophotometer at 595 nm, calibrated with a buffer and Bradford reagent-only blank. Each sample was measured three times and the mean value was plotted against a standard curve to extrapolate the total protein content in the sample. The standard curve was prepared using eight serial dilutions of bovine serum albumin (BSA, 1mg/ml, Sigma) between 0 and 50 µl diluted in buffer to a total volume of 1 ml. Protein content estimates were multiplied by 30 to express protein content per egg (µg).

To assess the impact of outgroup threat on parental care (Experiment I), we filmed (Sony Handycam HDR-XR520) each group for 10 min on the morning after a clutch was laid, before they experienced an intrusion. Parental care at the egg stage, consisting of fanning and cleaning the eggs, aids embryonic development and survival (23). From the videos, we recorded all clutch visits and parental-care behaviours displayed by all group members, as well as the total time spent within a body length of the clutch and in parental activities. Egg-cleaning and clutch-fanning were analysed both separately and together to assess overall parental care.

### Reproductive output

We visually counted the number of surviving offspring on the day that they reached 1-month post-hatching. To count offspring numbers in larger clutches reliably, we temporarily divided the tanks into three sections using transparent partitions and counted the number of offspring in each section separately. All counts were done twice to confirm totals; where values differed, we counted young a third time and either took the confirmed number of young or calculated the mean number from the three counts.

To assess the impacts of outgroup threat on offspring activity level and response to a sudden stimulus, we tested offspring at 1-month post-hatching. At this age, offspring actively explore the territory (i.e. swim around the tank), and when they perceive a threat, they sink to the substrate and remain immobile to blend with the sandy background (23,52). A sample of 5–10 offspring (N=21 clutches) was transferred from their home tank to a test container (20×20×10 cm), filled with 3 litres of water from their home tank, and left to settle for 30 min before the commencement of a trial. Each trial was filmed (Sony Handycam HDR-XR520) from above. After an initial 5-min undisturbed, pre-stimulus period, we released a small glass marble in a 60 cm plastic tube placed at a right angle relative to and touching the side of the container, so that the marble produced sudden vibrations and noise as it hit the container. We then filmed offspring behaviour for a 5 min post-stimulus period. We calculated mean offspring activity level pre-stimulus by observing 3 s of film every 20 s and recording how many offspring were actively swimming during that period. We also took screenshots every 20 s during the 5 min pre-stimulus period, from which we measured the mean nearest-neighbour distance of the offspring. From the post-stimulus period, we recorded the latency for all individuals to become immobile in response to the stimulus and the latency for the first offspring to become active again.

After the startle-stimulus trials, we euthanised the offspring with an overdose of tricaine methanesulfonate (MS222, 12 ml/ 100 ml tank water) and stored them in a 30% ethanol freshwater solution, shown to be adequate for simultaneously preserving body tissues and minimising shrinkage (77). We measured offspring standard length (from the tip of the snout to the end of the caudal peduncle, ±0.01 mm) using Leica Application Suite (version 4.4.0 (Build:454), Leica Microsystems Limited, Switzerland) connected to a camera (Greenough Stereozoom 0.8x manual) mounted on a stereo microscope (EZ4 HD, eyepiece 10x/21B). Immediately after measuring the offspring, we dried them for 36 h at 70°C before weighing them individually on a Mettler scale (AE260, Delta Range, ±0.1 mg). We then used the individual measurements to calculate mean offspring standard length and dry weight per clutch.

### Statistical analyses

All statistical analyses were conducted using RStudio (78) (version 1.2.5033). In Experiment I, we used paired t-tests or Wilcoxon signed-rank tests to assess intruder behaviour and defensive contributions between treatments and within individuals at the start and end of the study. We used a McNemar test and a Wilcoxon rank sum test to assess treatment differences in the likelihood of spawning and in the number of clutches produced, respectively. In Experiment II, we assessed treatment effects on clutch size with analysis of covariance (ANCOVA), which controlled for treatment duration and for female size. We analysed all other response variables in both experiments using linear mixed-effects models (LMMs) with the package “lme4” (79) (version 1.1-21). To simplify the mixed-effects models, we conducted stepwise backward elimination of terms starting with least significant interactions followed by least significant covariates and main factors. We assessed term significance by comparing a model with and without the specific term using likelihood ratio tests (chi-square tests using R function Anova (80)).

All mixed-effects models initially included clutch identity, focal group identity and tank-triplet identity as random factors to control for repeated observations from the same clutch and group and for the shared neighbours within each triplet, respectively. However, because in both experiments several groups did not spawn in the two treatments, we removed triplet identity from the analyses in models where it explained zero random variance. Our main factors of interest were treatment (Intruded and Control) and its interaction with treatment duration (covariate); we controlled for the effects of several covariates in different models as appropriate (full details of factors used in each analysis in tables of model outputs in the Supplementary Information). The effects of significant interactions between treatment and treatment duration were teased apart by analysing the effect of treatment duration on each treatment separately. We visually assessed the normality and homogeneity of residual distributions by inspecting qqplots and plots of residuals against fitted values. We then applied the “gaussianisation” algorithm in the “lambertW” package, based on Goerg (81), to normalise skewed residual distributions in the analyses of the following response variables: latency to spawn, hatching success, number of caring events towards a clutch, time spent cleaning and time spent caring for a clutch in Experiment I. Response variables in Experiment II did not require transformation. Analyses of hatching success were done on the subset of clutches that had some hatching success. Five clutches with zero hatching success (three Control clutches from the same group, two Intruded clutches from separate groups) were removed from the analysis because they were either eaten or failed to develop due to fungal infections; factors unrelated to the treatments. Significant parameter estimates and associated 95% confidence intervals or treatment effect sizes are provided in the text; all parameter estimates are provided in the Supplementary Information.

## Acknowledgments

We thank Richard Wall for access to his laboratory facilities, Shatha Alqurashi and Saeed Alasmari for assistance in developing protein and lipid extraction protocols, Martin Aveling for the artwork in Figure 1, Barbara Taborsky and Michael Taborsky for detailed discussions about this project and the study species across the years, and Stephanie King, Ben Ashton and Patrick Kennedy for comments on the manuscript. This work was supported by a European Research Council Consolidator Grant (project no. 682253) awarded to A.N.R.

## Author contributions

Both authors designed the experiments. I.B.G. performed the experiments and analyses. I.B.G. drafted the manuscript and both authors contributed to subsequent writing.

## Data availability

The data that support the findings of this study will be made available in the Dryad Digital Repository.

## Supplementary information

**Table S1.** Statistical summary of linear mixed models testing the effect of chronic outgroup conflict (Intruded vs Control, Experiment I) on breeding timings. Effect of outgroup conflict on (a) latency to spawn (days), and (b) inter-clutch interval (days); latency to spawn was transformed using a “gaussianisation” algorithm (81) to normalise the residual distributions. Female size relates to dominant female standard length at the start of the study. Group identity nested within tank-triplet identity or just tank-triplet identity were fitted as random intercepts (with standard deviations shown). The reference level for Treatment was Control. Non-significant fixed terms are shown by order of removal. Significant terms are shown in bold; near-significant terms in final model are shown in italics.

| FINAL MODEL | $X^2$ | t-value | d.f. | p | estimate | s.e. | C.I. |
| --- | --- | --- | --- | --- | --- | --- | --- |
| **Intercept** |  | **2.37** | **8.07** | **0.045** | **11.96** | **5.06** | **1.24–22.22** |
| REMOVED TERMS | $X^2$ | | d.f. | p | | | |
| Treatment x Female size | 0.45 |  | 1 | 0.502 |  |  |  |
| Treatment | 1.07 |  | 1 | 0.301 |  |  |  |
| Female size | 2.27 |  | 1 | 0.132 |  |  |  |

b. Inter-clutch interval (N=21 intervals) Tank-triplet/Group: 6.87; Tank-triplet: 6.01; Residual: 2.16
| FINAL MODEL | $X^2$ | t-value | d.f. | p | estimate | s.e. | C.I. |
| --- | --- | --- | --- | --- | --- | --- | --- |
| Intercept |  | -0.58 | 10.82 | 0.570 | -5.72 | 9.85 | -26.08–13.63 |
| **Treatment** | **3.89** |  | **1** | **0.049** |  |  |  |
| *Intruded* |  | *2.05* | *5.86* | *0.088* | *8.88* | *4.34* | *0.06–17.71* |
| **Female size** | **7.32** |  | **1** | **0.007** |  |  |  |
| ***Female size*** |  | **2.97** | **8.64** | **0.017** | **0.49** | **0.17** | **0.16–0.85** |
| REMOVED TERMS | $X^2$ | | d.f. | p | | | |
| Treatment x Female size | 0.80 |  | 1 | 0.373 |  |  |  |

**Table S2.**
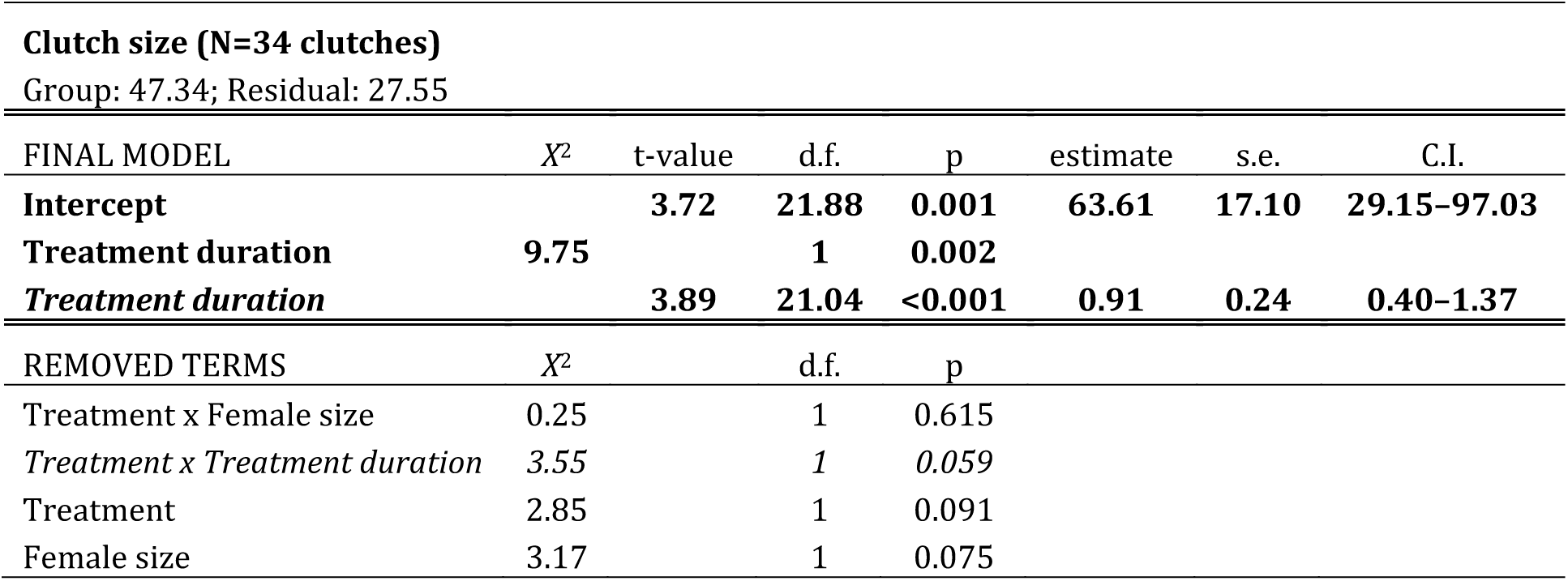
Statistical summary of a linear mixed model testing the effect of chronic outgroup conflict (Intruded vs Control, Experiment I) on clutch size. Female size relates to the standard-length measurement of the dominant female made closest in time to the production of each clutch (start or end of study). Group identity was fitted as a random intercept (with standard deviation shown). The reference level for Treatment was Control. Non-significant fixed terms are shown by order of removal. Significant terms are shown in bold; removed near-significant terms of interest are shown in italics.

**Table S3.**
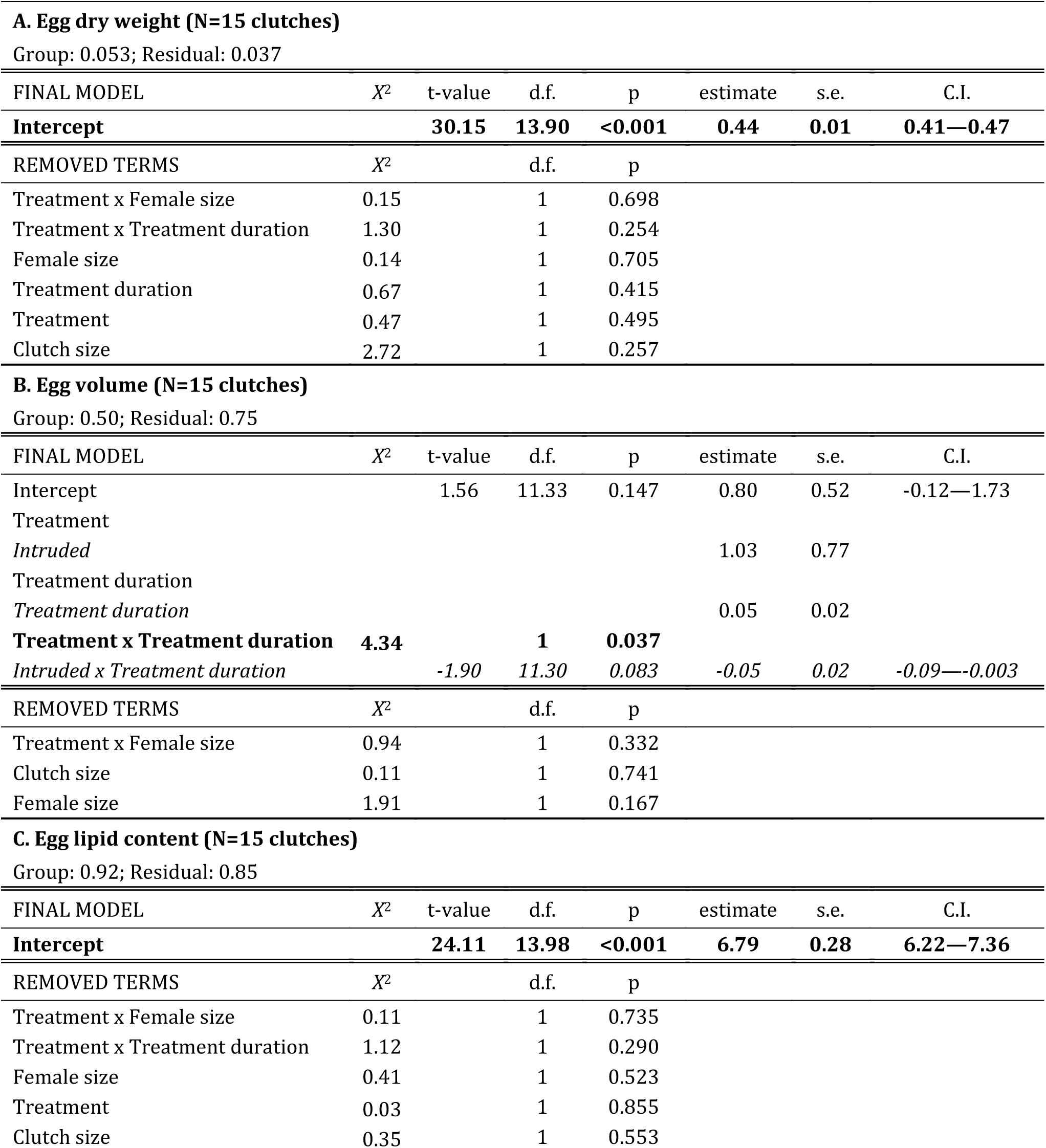
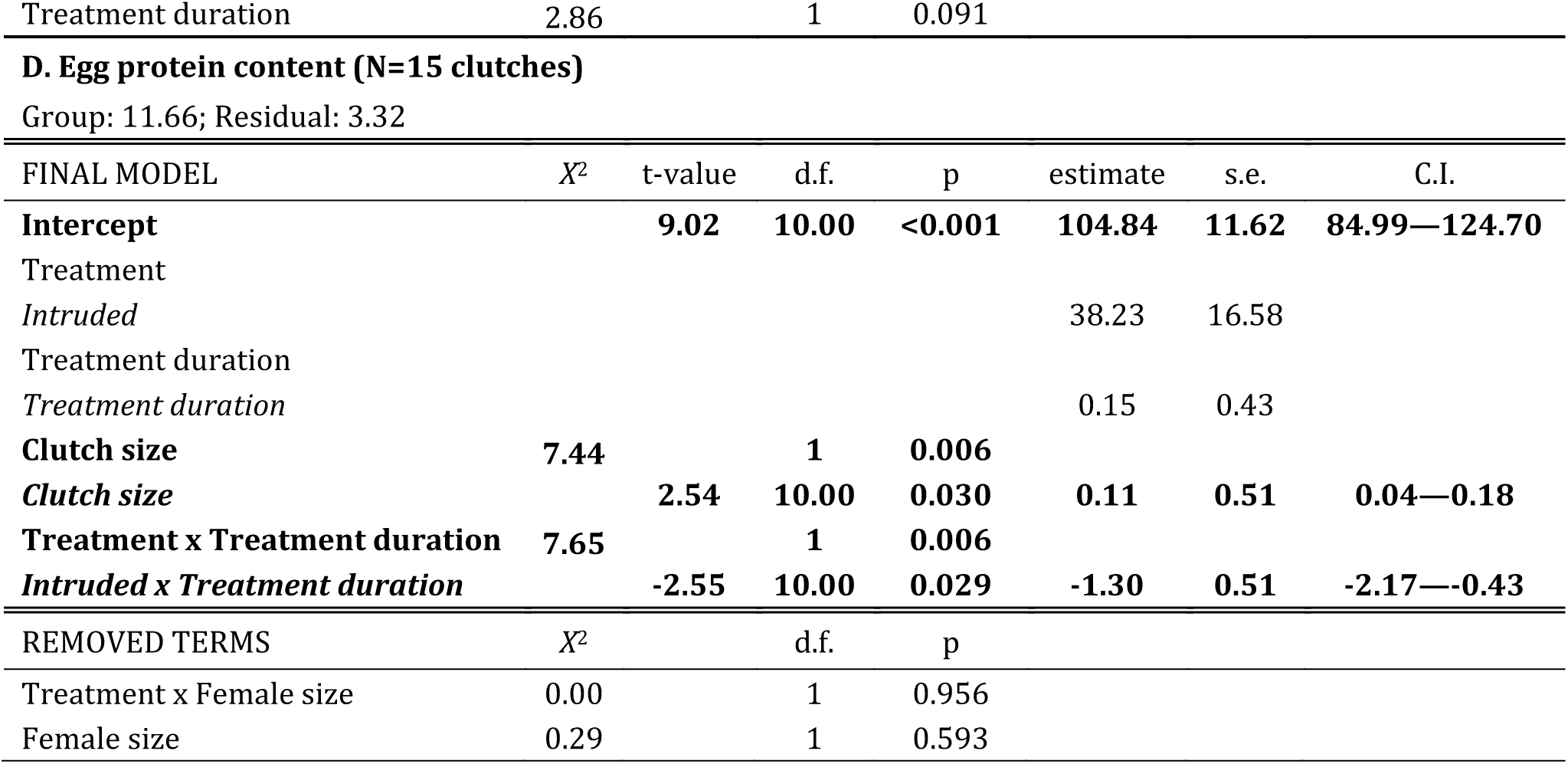
Statistical summary of linear mixed models testing the effect of chronic outgroup conflict (Intruded vs Control, Experiment II) on mean morphological and physiological egg characters. Effect of outgroup conflict on egg (a) volume (mm^3^), (b) dry weight (mg), (c) protein content (µg) and (d) lipid content (µg). Female size relates to dominant female standard length at the start of the study. Group identity was fitted as a random intercept (with standard deviation shown). The reference level for Treatment was Control. For fixed effects included in significant interactions, only parameter estimates are shown. Non-significant fixed terms are shown by order of removal. Significant terms are shown in bold, near-significant terms in final model are shown in italics.

**Table S4.**
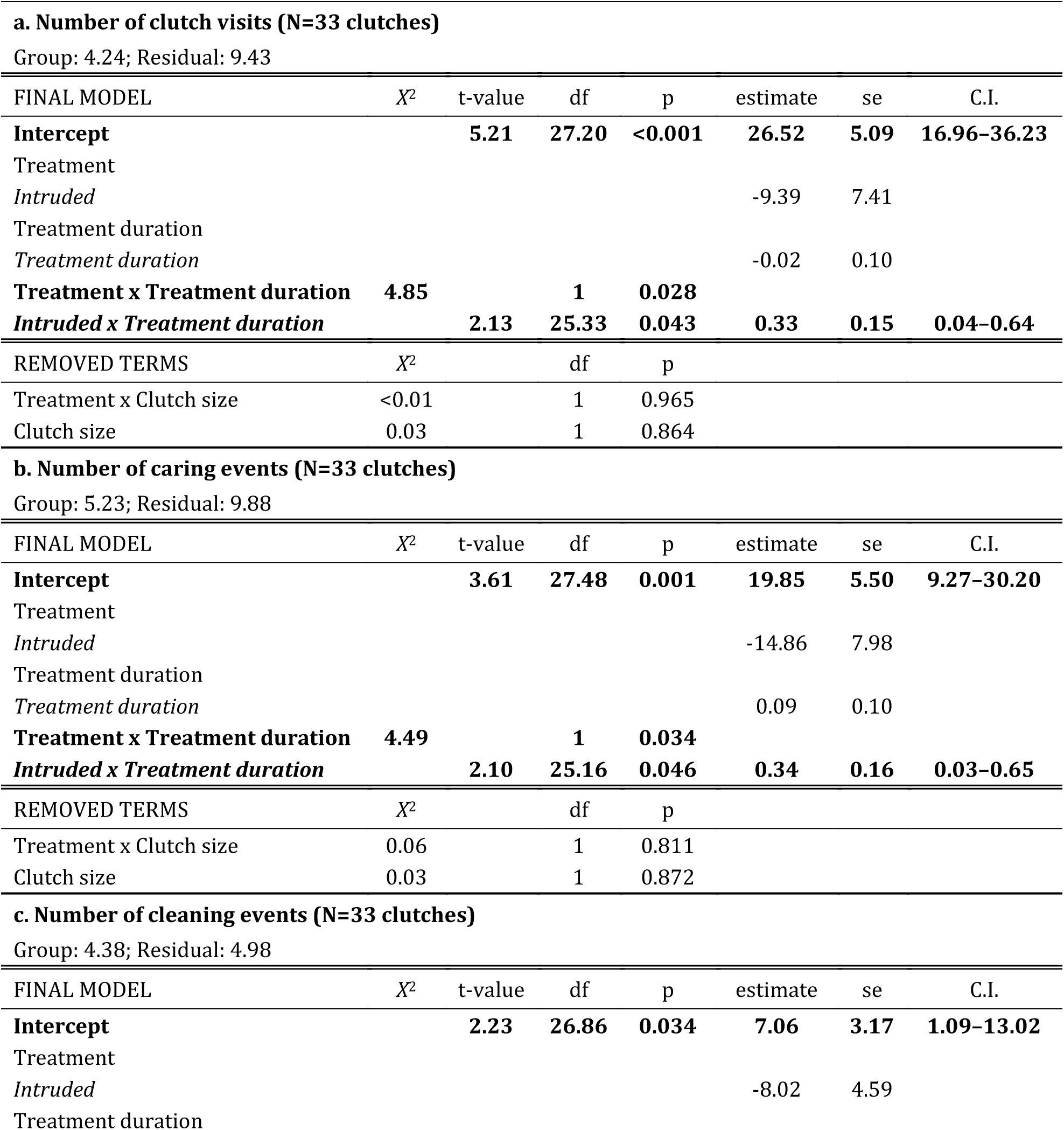
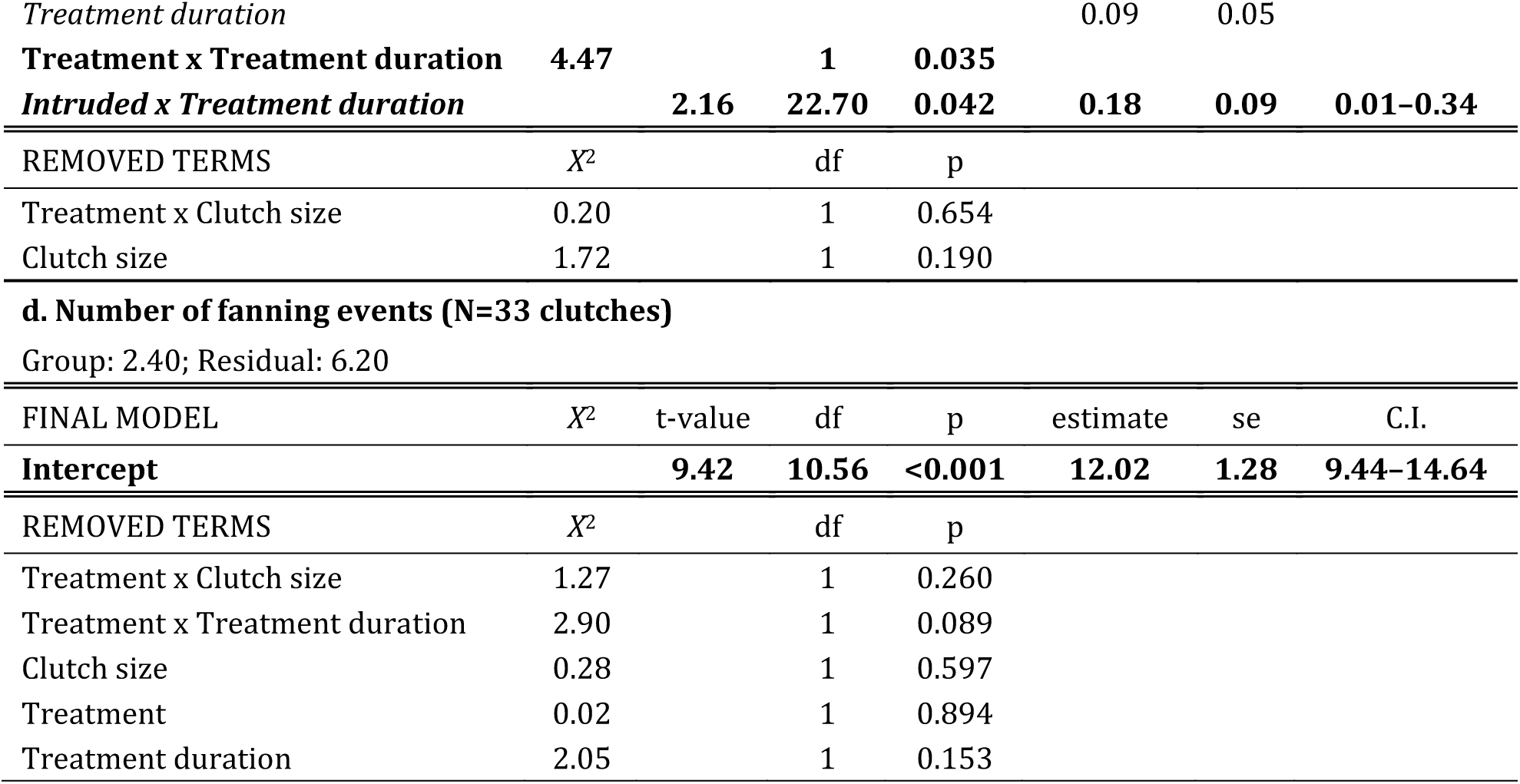
Statistical summary of linear mixed models testing the effect of chronic outgroup conflict (Intruded vs Control, Experiment I) on number of parental-care behaviours performed during a 10-min period. Effect of outgroup conflict on (a) clutch visits, (b) caring (egg-cleaning and fanning) acts, (c) egg-cleaning, and (d) egg-fanning; number of caring behaviours was transformed using a “gaussianisation” algorithm (81) to normalise the residual distributions. Group identity was fitted as a random intercept (with standard deviation shown). The reference level for Treatment was Control. For fixed effects included in significant interactions, only parameter estimates are shown. Non-significant fixed terms are shown by order of removal. Significant terms are shown in bold, near-significant terms in final models are shown in italics.

| **a. Number of clutch visits (N=33 clutches)** |  |  |  |  |  |  |  |
| --- | --- | --- | --- | --- | --- | --- | --- |
| Group: 4.24; Residual: 9.43 |  |  |  |  |  |  |  |
| FINAL MODEL | X<sup>2</sup> | t-value | df | p | estimate | se | C.I. |
| **Intercept** |  | **5.21** | **27.20** | **<0.001** | **26.52** | **5.09** | **16.96–36.23** |
| Treatment |  |  |  |  |  |  |  |
| *Intruded* |  |  |  |  | -9.39 | 7.41 |  |
| Treatment duration |  |  |  |  |  |  |  |
| *Treatment duration* |  |  |  |  | -0.02 | 0.10 |  |
| **Treatment x Treatment duration** | **4.85** |  | **1** | **0.028** |  |  |  |
| ***Intruded x Treatment duration*** |  | **2.13** | **25.33** | **0.043** | **0.33** | **0.15** | **0.04–0.64** |
| REMOVED TERMS | X<sup>2</sup> |  | df | p |  |  |  |
| Treatment x Clutch size | <0.01 |  | 1 | 0.965 |  |  |  |
| Clutch size | 0.03 |  | 1 | 0.864 |  |  |  |
| **b. Number of caring events (N=33 clutches)** |  |  |  |  |  |  |  |
| Group: 5.23; Residual: 9.88 |  |  |  |  |  |  |  |
| FINAL MODEL | X<sup>2</sup> | t-value | df | p | estimate | se | C.I. |
| **Intercept** |  | **3.61** | **27.48** | **0.001** | **19.85** | **5.50** | **9.27–30.20** |
| Treatment |  |  |  |  |  |  |  |
| *Intruded* |  |  |  |  | -14.86 | 7.98 |  |
| Treatment duration |  |  |  |  |  |  |  |
| *Treatment duration* |  |  |  |  | 0.09 | 0.10 |  |
| **Treatment x Treatment duration** | **4.49** |  | **1** | **0.034** |  |  |  |
| ***Intruded x Treatment duration*** |  | **2.10** | **25.16** | **0.046** | **0.34** | **0.16** | **0.03–0.65** |
| REMOVED TERMS | X<sup>2</sup> |  | df | p |  |  |  |
| Treatment x Clutch size | 0.06 |  | 1 | 0.811 |  |  |  |
| Clutch size | 0.03 |  | 1 | 0.872 |  |  |  |
| **c. Number of cleaning events (N=33 clutches)** |  |  |  |  |  |  |  |
| Group: 4.38; Residual: 4.98 |  |  |  |  |  |  |  |
| FINAL MODEL | X<sup>2</sup> | t-value | df | p | estimate | se | C.I. |
| **Intercept** |  | **2.23** | **26.86** | **0.034** | **7.06** | **3.17** | **1.09–13.02** |
| Treatment |  |  |  |  |  |  |  |
| *Intruded* |  |  |  |  | -8.02 | 4.59 |  |
| Treatment duration |  |  |  |  |  |  |  |
| *Treatment duration* |  |  |  |  | 0.09 | 0.05 |  |
| **Treatment x Treatment duration** | **4.47** |  | **1** | **0.035** |  |  |  |
| **Intruded x Treatment duration** |  | **2.16** | **22.70** | **0.042** | **0.18** | **0.09** | **0.01-0.34** |
| REMOVED TERMS | *X*<sup>2</sup> |  | df | p |  |  |  |
| Treatment x Clutch size | 0.20 |  | 1 | 0.654 |  |  |  |
| Clutch size | 1.72 |  | 1 | 0.190 |  |  |  |
| FINAL MODEL | *X*<sup>2</sup> | t-value | df | p | estimate | se | C.I. |
| **Intercept** |  | **9.42** | **10.56** | **<0.001** | **12.02** | **1.28** | **9.44-14.64** |
| REMOVED TERMS | *X*<sup>2</sup> |  | df | p |  |  |  |
| Treatment x Clutch size | 1.27 |  | 1 | 0.260 |  |  |  |
| Treatment x Treatment duration | 2.90 |  | 1 | 0.089 |  |  |  |
| Clutch size | 0.28 |  | 1 | 0.597 |  |  |  |
| Treatment | 0.02 |  | 1 | 0.894 |  |  |  |
| Treatment duration | 2.05 |  | 1 | 0.153 |  |  |  |

**Table S5.**
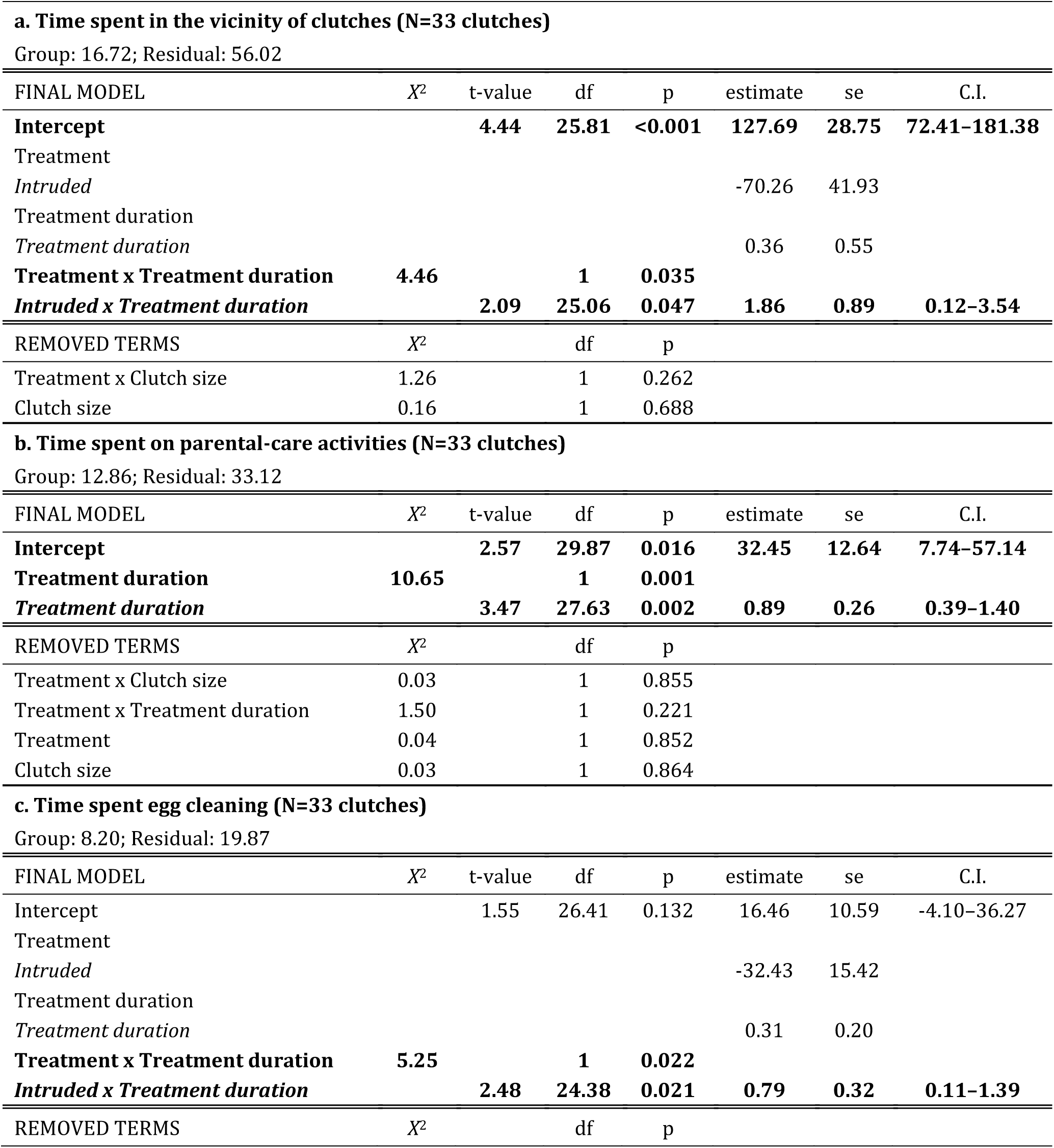
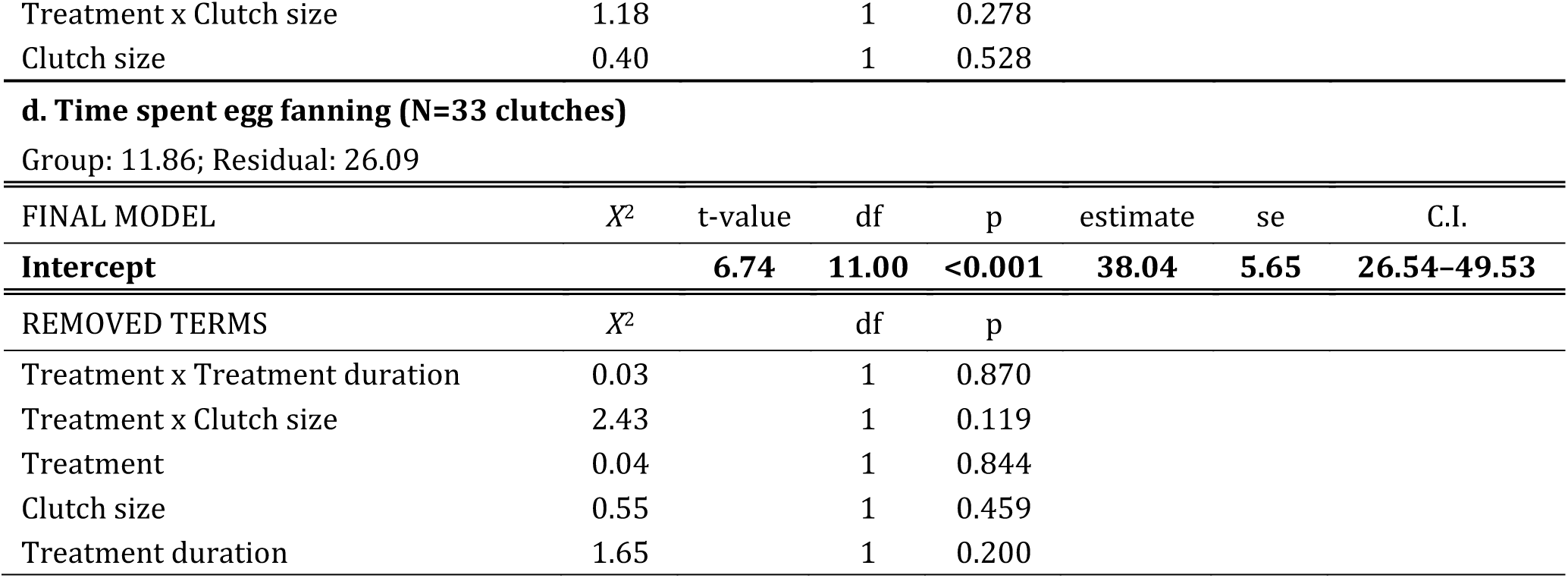
Statistical summary of linear mixed models testing the effect of chronic outgroup conflict (Intruded vs Control, Experiment I) on time (s) spent on parental-care behaviour during a 10-min period. Effect of outgroup conflict on time spent (a) in vicinity of clutch, (b) on parental-care activities (egg-cleaning and fanning), (c) on egg-cleaning, and (d) on egg-fanning; time spent in parental-care activities and egg-cleaning were transformed using a “gaussianisation” algorithm (81) to normalise the residual distributions. Group identity was fitted as a random intercept (with standard deviation shown). The reference level for Treatment was Control. For fixed effects included in significant interactions, only parameter estimates are shown. Non-significant fixed terms are shown by order of removal. Significant terms are shown in bold.

**Table S6.**
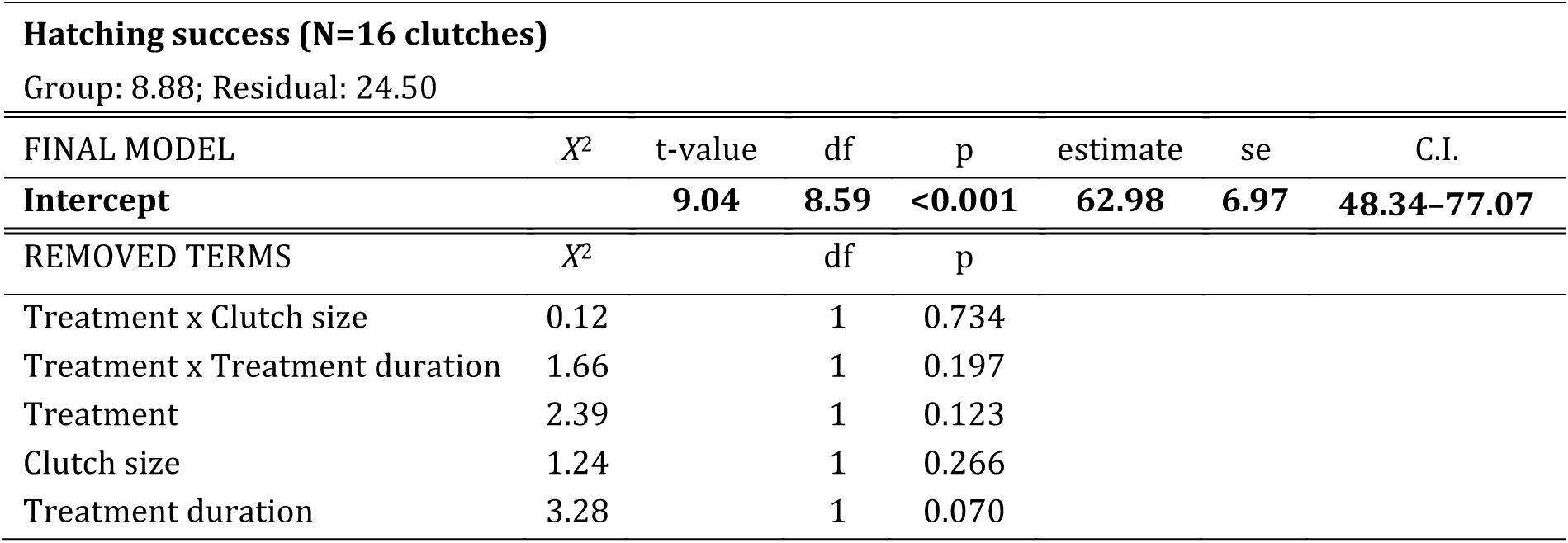
Statistical summary of a linear mixed model testing the effect of chronic outgroup conflict (Intruded vs Control, Experiment I) on hatching success (%). Hatching success was transformed using a “gaussianisation” algorithm (81) to normalise the residual distributions. Group identity was fitted as a random intercept (with standard deviation shown). The reference level for Treatment was Control. Non-significant fixed terms are shown by order of removal. Significant term is shown in bold.

**Table S7.** Statistical summary of linear mixed models testing the effect of chronic outgroup conflict (Intruded vs Control, Experiment I) on offspring survival to 1-month post-hatching. Effect of outgroup conflict on (a) number of surviving offspring, (b) relative survival from hatching (%) and (c) number of surviving offspring from hatching. Group identity was fitted as a random intercept (with standard deviation shown). The reference level for Treatment was Control. For fixed effects included in significant interactions, only parameter estimates are shown. Non-significant fixed terms are shown by order of removal. Significant terms are shown in bold.

| **a. Number of offspring surviving to 1-month of age (N=27 clutches)** |  |  |  |  |  |  |  |
| --- | --- | --- | --- | --- | --- | --- | --- |
| Group: 4.76; Residual: 19.97 |  |  |  |  |  |  |  |
| FINAL MODEL | $X^2$ | t-value | df | p | estimate | se | C.I. |
| Intercept |  | -0.57 | 22.55 | 0.580 | -11.08 | 19.56 | -48.15–25.60 |
| Treatment |  |  |  |  |  |  |  |
| *Intruded* |  |  |  |  | 67.72 | 26.82 |  |
| Treatment duration |  |  |  |  | 0.49 | 0.27 |  |
| **Treatment x Treatment duration** | **6.58** |  | **1** | **0.010** |  |  |  |
| ***Intruded x Treatment duration*** |  | **-2.52** | **19.56** | **0.020** | **-0.96** | **0.38** | **-1.70– -0.25** |
| **b. Relative offspring survival from hatching to 1-month of age (N=15 clutches)** |  |  |  |  |  |  |  |
| Group: 26.59; Residual: 21.89 |  |  |  |  |  |  |  |
| FINAL MODEL | $X^2$ | t-value | df | p | estimate | se | C.I. |
| **Intercept** |  | **4.87** | **8.80** | **<0.001** | **52.44** | **10.76** | **30.69–75.16** |
| REMOVED TERMS | $X^2$ | | df | p | | | |
| Treatment x Treatment duration | 0.35 |  | 1 | 0.554 |  |  |  |
| Treatment x Number of hatchlings | 1.21 |  | 1 | 0.271 |  |  |  |
| Treatment duration | 0.04 |  | 1 | 0.842 |  |  |  |
| Treatment | 1.53 |  | 1 | 0.216 |  |  |  |
| Number of hatchlings | 1.37 |  | 1 | 0.243 |  |  |  |
| **c. Number of offspring surviving from hatching to 1-month of age (N=15 clutches)** |  |  |  |  |  |  |  |
| Group: 11.43; Residual: 11.20 |  |  |  |  |  |  |  |
| FINAL MODEL | $X^2$ | t-value | df | p | estimate | se | C.I. |
| Intercept |  | -1.04 | 11.00 | 0.322 | -11.86 | 11.44 | -33.45–8.42 |
| Treatment |  |  |  |  |  |  |  |
| *Intruded* |  |  |  |  | 15.84 | 16.92 |  |
| Number of hatchlings |  |  |  |  | 0.90 | 0.19 |  |
| **Treatment x Number of hatchlings** | **5.12** |  | **1** | **0.024** |  |  |  |
| ***Intruded x Number of hatchlings*** |  | **-2.47** | **10.02** | **0.033** | **-0.65** | **0.26** | **-1.11– -0.12** |
| REMOVED TERMS | $X^2$ | | df | p | | | |
| Treatment x Treatment duration | 0.32 |  | 1 | 0.571 |  |  |  |
| Treatment duration | 0.13 |  | 1 | 0.721 |  |  |  |

**Table S8.** Statistical summary of linear mixed models testing the effect of chronic outgroup conflict (Intruded vs Control, Experiment I) on offspring behaviour. Effect of outgroup conflict on (a) mean pre-stimulus activity (%), (b) mean pre-stimulus nearest-neighbour distance (cm) (c) mean latency to freeze (s) post-stimulus, and (d) latency (s) to become active post-stimulus. Group identity was fitted as a random intercept (with standard deviation shown). The reference level for Treatment was Control. For fixed effects included in significant interactions, only parameter estimates are shown. Non-significant fixed terms are shown by order of removal. Significant terms are shown in bold, near-significant terms in final models are shown in italics.

| **a. Pre-stimulus activity (N=21 clutches)** |  |  |  |  |  |  |  |
| --- | --- | --- | --- | --- | --- | --- | --- |
| Group: 0.11; Residual: 0.17 |  |  |  |  |  |  |  |
| FINAL MODEL | $X^2$ | t-value | df | p | estimate | se | C.I. |
| **Intercept** |  | **8.02** | **15.88** | **<0.001** | **1.09** | **0.14** | **0.77–1.36** |
| **Treatment duration** | **5.40** |  | **1** | **0.020** |  |  |  |
| ***Treatment duration*** |  | **-2.81** | **13.16** | **0.015** | **-0.01** | **0.00** | **-0.009– -0.001** |
| REMOVED TERMS | $X^2$ | | df | p | | | |
| Treatment x Treatment duration | 1.41 |  | 1 | 0.234 |  |  |  |
| Offspring size | 0.28 |  | 1 | 0.596 |  |  |  |
| Number of offspring | 0.35 |  | 1 | 0.557 |  |  |  |
| Treatment | 0.95 |  | 1 | 0.329 |  |  |  |
| **b. Nearest-neighbour distance (N=21 clutches)** |  |  |  |  |  |  |  |
| Group: 0.00; Residual: 0.61 |  |  |  |  |  |  |  |
| FINAL MODEL | $X^2$ | t-value | df | p | estimate | se | C.I. |
| **Intercept** |  | **6.13** | **19.00** | **<0.001** | **6.93** | **1.13** | **4.72–9.13** |
| **Number of offspring** | **11.18** |  | **1** | **<0.001** |  |  |  |
| ***Number of offspring*** |  | **-3.65** | **19.00** | **0.002** | **-0.43** | **0.12** | **-0.65– -0.20** |
| REMOVED TERMS | $X^2$ | | df | p | | | |
| Treatment x Treatment duration | 2.14 |  | 1 | 0.143 |  |  |  |
| Treatment | 0.13 |  | 1 | 0.714 |  |  |  |
| Offspring size | 0.15 |  | 1 | 0.701 |  |  |  |
| Treatment duration | 0.94 |  | 1 | 0.333 |  |  |  |
| **c. Latency to freeze (N=21 clutches)** |  |  |  |  |  |  |  |
| Group: 0.70; Residual: 0.68 |  |  |  |  |  |  |  |
| FINAL MODEL | $X^2$ | t-value | df | p | estimate | se | C.I. |
| *Intercept* |  | *-2.07* | *17.62* | *0.054* | *-2.74* | *1.33* | *-5.50 –0.12* |
| **Treatment duration** | **6.94** |  | **1** | **0.008** |  |  |  |
| ***Treatment duration*** |  | **3.20** | **14.43** | **0.006** | **0.03** | **0.01** | **0.01–0.06** |
| **Pre-stimulus activity levels** | **11.90** |  | **1** | **<0.001** |  |  |  |
| ***Pre-stimulus activity levels*** |  | **4.16** | **16.33** | **<0.001** | **4.38** | **1.05** | **2.05–6.52** |
| REMOVED TERMS | $X^2$ | | df | p | | | |
| Treatment x Treatment duration | 0.64 |  | 1 | 0.424 |  |  |  |
| Offspring size | 0.01 |  | 1 | 0.931 |  |  |  |
| Treatment | 0.03 |  | 1 | 0.855 |  |  |  |
| Number of offspring | 3.74 |  | 1 | 0.053 |  |  |  |

**d. Latency to first movement post-stimulus (N=21 clutches)** Group: 0.00 ± 0.00; Residual: 155.00 ± 12.45
| FINAL MODEL | $X^2$ | t-value | df | p | estimate | se | C.I. |
| --- | --- | --- | --- | --- | --- | --- | --- |
| Intercept |  | 1.68 | 16.00 | 0.112 | 35.15 | 20.87 | -2.25 -72.55 |
| Treatment |  |  |  |  |  |  |  |
| *Intruded* |  |  |  |  | 34.22 | 17.87 |  |
| Treatment duration |  |  |  |  |  |  |  |
| *Treatment duration* |  |  |  |  | 0.11 | 0.22 |  |
| **Pre-stimulus activity levels** | **3.88** |  | **1** | **0.049** |  |  |  |
| *Pre-stimulus activity levels* |  | -1.80 | 16.00 | 0.091 | -26.12 | 14.51 | -52.12 - -0.13 |
| **Treatment x Treatment duration** | **4.62** |  | **1** | **0.032** |  |  |  |
| *Treatment x Treatment duration* |  | -2.04 | 16.00 | 0.059 | -0.55 | 0.27 | -1.03 - -0.07 |
| REMOVED TERMS | $X^2$ | | df | p | | | |
| Number of offspring | 1.43 |  | 1 | 0.232 |  |  |  |
| Offspring size | 3.04 |  | 1 | 0.081 |  |  |  |
| Number of offspring | 1.52 |  | 1 | 0.217 |  |  |  |

**Table S9.** Statistical summary of linear mixed models testing the effect of chronic outgroup conflict (Intruded vs Control, Experiment I) on offspring size. Effect of outgroup size on (a) mean standard length (mm) and (b) mean dry weight (mg). Group identity nested within tank-triplet identity or just group identity were fitted as random intercepts (with standard deviations shown). The reference level for Treatment was Control. Non-significant fixed terms are shown by order of removal. Significant terms are shown in bold, removed near-significant terms of interest are shown in italics.

| **a. Offspring standard length (N=24 clutches)** |  |  |  |  |  |  |  |
| --- | --- | --- | --- | --- | --- | --- | --- |
| Group: 1.22; Residual: 0.61 |  |  |  |  |  |  |  |
| FINAL MODEL | X<sup>2</sup> | t-value | df | p | estimate | se | C.I. |
| **Intercept** |  | **16.05** | **20.34** | **<0.001** | **11.71** | **0.73** | **10.26–13.11** |
| **Treatment** | **4.89** |  | **1** | **0.027** |  |  |  |
| *Intruded* |  | **-2.25** | **9.62** | **0.049** | **-1.68** | **0.75** | **-3.13– -0.23** |
| **Treatment duration** | **9.72** |  | **1** | **0.002** |  |  |  |
| *Treatment duration* |  | **-4.36** | **12.42** | **<0.001** | **-0.03** | **0.01** | **-0.05– -0.02** |
| REMOVED TERMS | X<sup>2</sup> |  | df | p |  |  |  |
| Treatment x Treatment duration | 2.15 |  | 1 | 0.143 |  |  |  |
| Number of living offspring | 3.30 |  | 1 | 0.069 |  |  |  |
| **b. Offspring dry weight (N=24 clutches)** |  |  |  |  |  |  |  |
| Tank-triplet/Group: 1.01; Tank-triplet: 1.47; Residual: 1.20 |  |  |  |  |  |  |  |
| FINAL MODEL | X<sup>2</sup> | t-value | df | p | estimate | se | C.I. |
| **Intercept** |  | **6.73** | **6.84** | **<0.001** | **4.23** | **0.63** | **2.95–5.56** |
| REMOVED TERMS | X<sup>2</sup> |  | df | p |  |  |  |
| Treatment x Treatment duration | 0.15 |  | 1 | 0.695 |  |  |  |
| Treatment duration | 2.56 |  | 1 | 0.110 |  |  |  |
| Number of living offspring | 1.74 |  | 1 | 0.187 |  |  |  |
| *Treatment* | *3.15* |  | *1* | *0.076* |  |  |  |

